# The brainstem’s red nucleus was evolutionarily upgraded to support goal-directed action

**DOI:** 10.1101/2023.12.30.573730

**Authors:** Samuel R. Krimmel, Timothy O. Laumann, Roselyne J. Chauvin, Tamara Hershey, Jarod L. Roland, Joshua S. Shimony, Jon T. Willie, Scott A. Norris, Scott Marek, Andrew N. Van, Julia Monk, Kristen M. Scheidter, Forrest Whiting, Nadeshka Ramirez-Perez, Athanasia Metoki, Anxu Wang, Benjamin P. Kay, Hadas Nahman-Averbuch, Damien A. Fair, Charles J. Lynch, Marcus E. Raichle, Evan M. Gordon, Nico U.F. Dosenbach

**Affiliations:** Department of Neurology, Washington University School of Medicine, St. Louis, Missouri, USA; Department of Psychiatry, Washington University School of Medicine, St. Louis, Missouri, USA; Mallinckrodt Institute of Radiology, Washington University School of Medicine, St. Louis, Missouri, USA; Department of Psychological & Brain Sciences, Washington University, St. Louis, Missouri, USA; Department of Neurosurgery, Washington University School of Medicine, St. Louis, Missouri, USA; Department of Psychiatry, Weill Cornell Medicine, New York, New York, USA; Department of Neuroscience, Washington University School of Medicine, St. Louis, Missouri; Department of Biomedical Engineering, Washington University, St. Louis, Missouri; Division of Computation and Data Science, Washington University, St. Louis, Missouri, USA; Washington University Pain Center, Department of Anesthesiology, Washington University School of Medicine, St. Louis, Missouri, USA; Department of Pediatrics, University of Minnesota, Minneapolis, Minnesota, USA; Masonic Institute for the Developing Brain, University of Minnesota, Minneapolis, Minnesota, USA; Institute of Child Development, University of Minnesota, Minneapolis, Minnesota, USA; Program in Occupational Therapy, Washington University, St. Louis, Missouri, USA; Department of Pediatrics, Washington University School of Medicine, St. Louis, Missouri, USA

**Author notes:** **Declaration of Competing Interest** D.A.F., and N.U.F.D. have a financial interest in Turing Medical Inc. and may benefit financially if the company is successful in marketing FIRMM motion monitoring software products. A.N.V., D.A.F. and N.U.F.D. may receive royalty income based on FIRMM technology developed at Washington University School of Medicine and Oregon Health and Sciences University and licensed to Turing Medical Inc. D.A.F. and N.U.F.D. are co-founders of Turing Medical Inc. These potential conflicts of interest have been reviewed and are managed by Washington University School of Medicine, Oregon Health and Sciences University and the University of Minnesota. The other authors declare no competing interests.

## Abstract

The red nucleus is a large brainstem structure that coordinates limb movement for locomotion in quadrupedal animals (Basile et al., 2021). The humans red nucleus has a different pattern of anatomical connectivity compared to quadrupeds, suggesting a unique purpose (Hatschek, 1907). Previously the function of the human red nucleus remained unclear at least partly due to methodological limitations with brainstem functional neuroimaging (Sclocco et al., 2018). Here, we used our most advanced resting-state functional connectivity (RSFC) based precision functional mapping (PFM) in highly sampled individuals (n = 5) and large group-averaged datasets (combined *N* ∼ 45,000), to precisely examine red nucleus functional connectivity.

Notably, red nucleus functional connectivity to motor-effector networks (somatomotor hand, foot, and mouth) was minimal. Instead, red nucleus functional connectivity along the central sulcus was specific to regions of the recently discovered somato-cognitive action network (SCAN; (Gordon et al., 2023)). Outside of primary motor cortex, red nucleus connectivity was strongest to the cingulo-opercular (CON) and salience networks, involved in action/cognitive control (Dosenbach et al., 2007; Newbold et al., 2021) and reward/motivated behavior (Seeley, 2019), respectively. Functional connectivity to these two networks was organized into discrete dorsal-medial and ventral-lateral zones. Red nucleus functional connectivity to the thalamus recapitulated known structural connectivity of the dento-rubral thalamic tract (DRTT) and could prove clinically useful in functionally targeting the ventral intermediate (VIM) nucleus. In total, our results indicate that far from being a ‘motor’ structure, the red nucleus is better understood as a brainstem nucleus for implementing goal-directed behavior, integrating behavioral valence and action plans.

## Introduction

The brainstem has been thought of an evolutionarily conserved structure, limited to physiological (e.g., breathing) and basic motor functions (e.g., locomotion) (Baizer, 2014), with the exception of neuromodulatory nuclei (e.g., locus coeruleus). The red nucleus, located in the midbrain, first emerged as quadruped precursors began coordinating extremities for movement (Basile et al., 2021; De Lange, 1912; Padel, 1993; Padel et al., 1986; Ten Donkelaar, 1988). The red nucleus contains magno- and parvo-cellular neurons (Basile et al., 2021). In quadrupeds, magnocellular red nucleus neurons project down the full length of the spinal cord, forming the rubrospinal tract, which evokes limb movements when stimulated (De Lange, 1912; Ghez, 1975). Parvocellular red nucleus neurons are smaller in diameter and do not project to the spinal cord (Basile et al., 2021). Instead, these neurons participate in the dento-rubro-thalamic tract (DRTT), which connects to the cerebellum’s dentate nucleus and the ventral intermediate nucleus (VIM) of the thalamus (Basile et al., 2021; Cacciola et al., 2019; Lapresle & Hamida, 1970). Due to the parvocellular red nucleus projection to the thalamus, clinical neuroscience primarily studies structural connectivity of the red nucleus as a tool to identify the VIM (Nowacki et al., 2022), which is a target for deep brain stimulation treatment of essential tremor and tremor predominant Parkinson’s Disease (Haubenberger & Hallett, 2018; Schlaier et al., 2015).

In a striking example of phylogenetic refinement from quadrupeds to bipedal humans, the proportion of red nucleus neurons has shifted strongly from magnocellular to parvocellular (Cisek, 2019; Massion, 1967; Padel et al., 1981). For instance, the reptilian red nucleus is almost entirely magnocellular (Massion, 1967), the feline red nucleus is approximately 2/3 magnocellular (Huisman et al., 1982), and the primate red nucleus is primarily parvocellular (Basile et al., 2021). Furthermore, comparison of the quadrupedal baboon to the bipedal upright gibbon shows that bipedalism coincides with a continued reduction of the rubrospinal tract (Padel et al., 1981). In humans, there is a small rubrospinal tract that only projects to the cervical spinal cord, indicating that it could only serve a minimal role in locomotion (Massion, 1988; Nathan & Smith, 1982). The proportion of cell types in the human red nucleus favors parvocellular so much so that studies of the red nucleus in humans are effectively studies of the parvocellular red nucleus (Nathan & Smith, 1982). Even though human locomotion is supported by the cortico-spinal tract rather than the rubro-spinal tract, the expansion of parvocellular neurons has maintained the red nucleus as the largest nucleus in the human midbrain.

Despite nearly 150 years of research, the red nucleus’s function in humans remains unclear (Basile et al., 2021), revealing major gaps in our understanding of a clinically relevant structure and the brainstem as a whole. Direct recordings from the parvocellular red nucleus show activity is unrelated to free-form movement in non-human primates (Herter et al., 2015; Kennedy et al., 1986) and humans (Lefranc et al., 2014). Interestingly, there appears to be a relationship between parvocellular red nucleus and goal-directed actions and cognition. In an arm fixation-maintenance study in non-human primates, arm fixation evoked no red nucleus response except when an adaptive arm correction was required (Herter et al., 2015). Human task fMRI indicates simple sensory stimulation and hand movements produce small red nucleus task activations relative to cognitive tactile discrimination tasks (Liu et al., 2000; Sung et al., 2022), and the human red nucleus may be active during cognitive control (de Hollander et al., 2017; Sung et al., 2022). Rodent electrophysiology recordings during a stop-signal task found trial-to-trial adjustments in the parvocellular red nucleus firing rate that were correlated with movement accuracy and speed, indicating proactive control signals in the red nucleus (Brockett et al., 2020). Based on these findings, some have argued for parvocellular red nucleus involvement in motor control (Basile et al., 2021), which is a broad concept including motor planning, execution, and feedback (Craighero et al., 1999; Fajen, 2009; Kalaska, 2009; Latash, 2012; Stanley & Miall, 2009; Taylor & Gottlieb, 1985; Vogt et al., 2003). Owing in part to structural connectivity to primary motor cortex, the adaptive control responses in parvocellular red nucleus could support a mechanism for indirect control of movements based on task goals, but the support for this hypothesis is weak.

Resting state functional connectivity (RSFC) has greatly expanded our understanding of human brain organization by revealing its subdivision into large-scale functional networks related to specific functions such as action control, movement and salience. (Biswal et al., 1995; Dosenbach et al., 2007; Power et al., 2011; Seeley et al., 2007; Yeo et al., 2011). With large amounts of high quality data it is now possible to identify networks at the individual level, a technique we have dubbed precision functional mapping (PFM (E. J. Allen et al., 2022; Gordon et al., 2017; Laumann et al., 2015; Lynch et al., 2020)). Using PFM, we previously mapped the functional connectivity profiles of the thalamus (Greene et al., 2020), cerebellum (Marek et al., 2018), and hippocampus (Zheng et al., 2021). This process has allowed us to test theories of subcortical nuclei, especially when such models argue for connectivity with specific networks. Unfortunately, brainstem functional neuroimaging has been historically limited due to low signal-to-noise ratio (SNR) related to head coil distance, suboptimal echo times, and unique forms of noise owing to cerebrospinal fluid pulsations (Beissner, 2015; de Hollander et al., 2015, 2017; Wright & Wald, 1997). As a result, fMRI and RSFC of the red nucleus have greatly lagged the rest of the brain, making it difficult to examine its organization in humans.

We have recently shown that the precentral gyrus (i.e. primary motor cortex) is separated into motor-effector specific regions (foot, hand, and mouth) and somato-cognitive action network (SCAN) regions for integrating body movement, goals, and physiology (Gordon et al., 2023). These SCAN regions are most closely coupled to the cingulo-opercular network (CON), which is important for executive action control (Dosenbach et al., 2007; Newbold et al., 2021). While the parvocellular red nucleus has structural connectivity with the primary motor cortex (Burman et al., 2000), it is unclear if this connectivity is with motor-effector or SCAN regions, which would have a fundamentally different interpretation. Should the red nucleus indirectly control movements based on task goals, we predict extensive connectivity with motor-effector networks in the primary motor cortex (i.e. somatomotor hand, food, and/or mouth). Importantly, parvocellular red nucleus structural connectivity extends far beyond the precentral gyrus, including a robust connection with the anterior cingulate cortex (Burman et al., 2000). The anterior cingulate contains many networks, including a large representation of the salience network, which is important for processing reward signals and motivation (Peters et al., 2016; Seeley, 2019; Seeley et al., 2007). Based on structural connectivity alone, it remains unknown which networks the human red nucleus is functionally connected with.

Here, we determined individual-specific RSFC of the human red nucleus by overcoming historically low signal-to-noise with novel denoising approaches. We verified PFM results using group-averaged data from three large fMRI datasets (Human Connectome Project (HCP), Adolescent Brain Cognitive Development (ABCD) study, UK Biobank (UKB); combined sample size of nearly 45,000 participants).

## Results

### The red nucleus is connected to salience and action control networks

Red nucleus (Fig 1A) functional connectivity was strongest in the dorsal anterior cingulate, medial prefrontal, pre-supplementary motor, insula (especially anterior insula), parietal operculum and anterior prefrontal cortex (Fig 1B, C). Functional connectivity was clearly organized into networks, with the strongest connectivity to the CON (action control; dorsal anterior cingulate cortex, anterior prefrontal cortex, and anterior insula (Dosenbach et al., 2007)), and salience network (reward/motivated behavior; anterior cingulate/medial prefrontal cortex and ventral anterior insula (Seeley et al., 2007)), but not to foot/hand/mouth effector-specific motor regions near the central sulcus (Fig 1B, C; SFig 1:4). Functional connectivity in the central sulcus was strongest to the SCAN regions, which are closely related to the CON (Gordon et al., 2023). Red nucleus was not connected to the default mode network regions in prefrontal cortex and fronto-parietal network regions in the lateral prefrontal cortex and insula. There were no obvious and consistent differences between the left and right red nucleus (SFig 1; SFig 3), nor were the results contingent on the functional connectivity threshold (SFig 2). This connectivity pattern was also evident in the three large group-averaged datasets totaling ∼ 45,000 participants (Fig 1C; SFig 1) and in additional individual-specific red nucleus seed maps (SFig 4).

**Figure 1:**
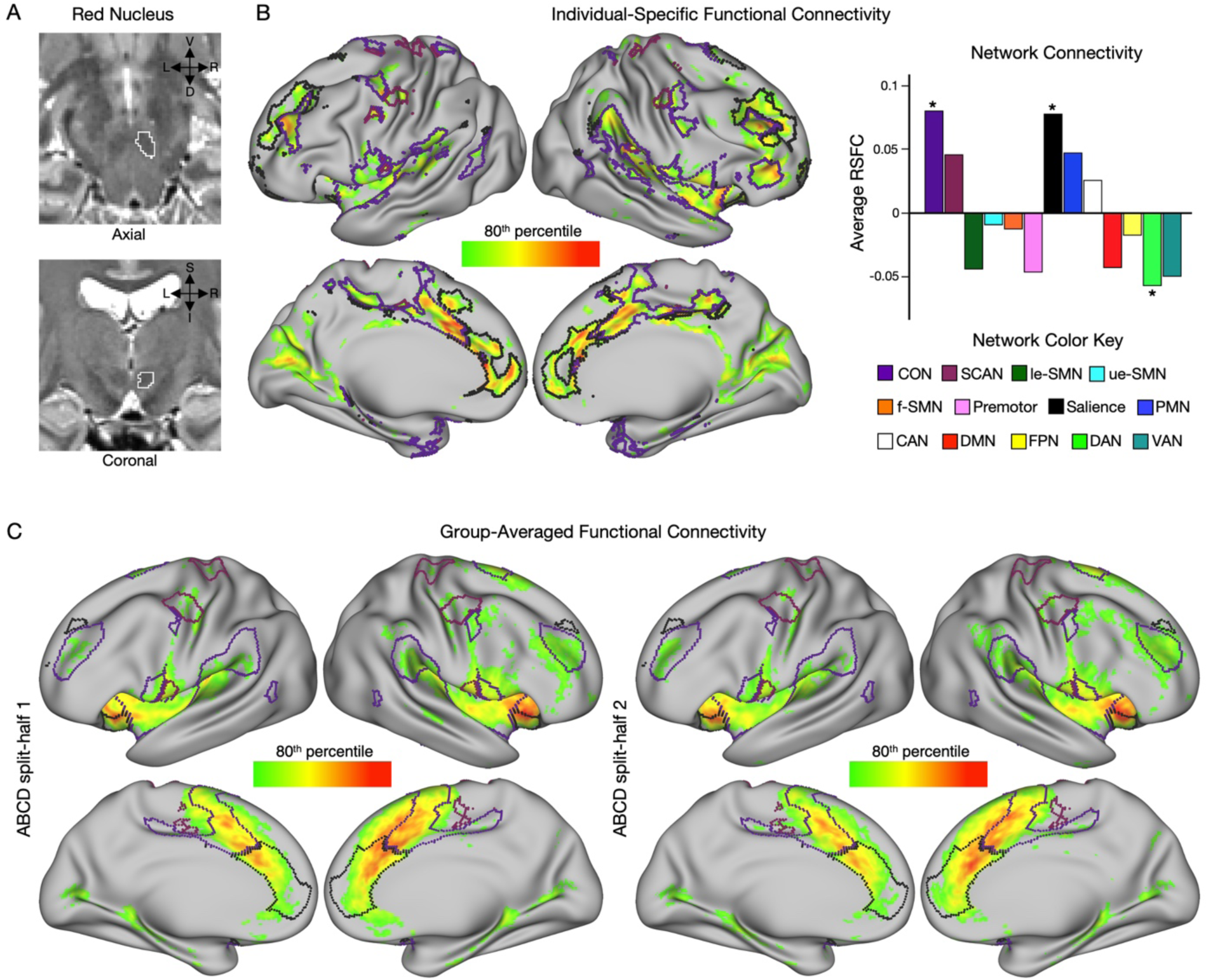
Functional connectivity mapping of the red nucleus. A) Axial (top) and coronal (bottom) display of the right red nucleus (white outline) overlaid on a T2w structural image for subject PFM-Nico. B) Resting state functional connectivity (RSFC) seeded from the right red nucleus in an exemplar highly sampled participant with multi-echo independent component analysis (MEICA) denoising (PFM-Nico; 134 min resting-state fMRI). Individual specific functional connectivity map shows strongest 20 percent of cortical vertices. Bar graph quantifies the average connectivity per network. The average connectivity was significantly different from zero for the salience, cingulo-opercular (CON) and dorsal attention (DAN) networks (two-sided t-test against null distribution, *P < 0.05, Bonferroni correction), but was only positive for salience and CON. C) Group-averaged functional connectivity map shows strongest 20 percent of cortical vertices using previously defined split-halves (Marek et al., 2022; n = 1964 participants each) from the Adolescent Brain Cognitive Development (ABCD) study. For additional participants see Supplemental Figures 1:4.

### Red nucleus is functionally connected to the ventral intermediate thalamus

Since the red nucleus is a node in the DRTT, we next examined connectivity to subcortical structures. Within thalamus, red nucleus functional connectivity was strongest to the ventral lateral posterior (VLP) nucleus, centered on the ventral intermediate nucleus (VIM), which is a major target for deep brain stimulation with a known structural connection to the red nucleus (Schlaier et al., 2015). This was observed at the individual level using a subject specific thalamic segmentation (Fig 2A) and verified using large group-averaged datasets (Fig 2B-D; UKB (n = 4,000), ABCD study (n = 3,928), HCP (n = 812)) and additional individual-specific analyses (SFig 5). Interestingly, red nucleus connectivity tended to be stronger in more dorsal sections of the VLP (MNI z coord. > 2 mm). Within cerebellum, red nucleus functional connectivity was primarily to lobule VI (SFig 6).

**Figure 2:**
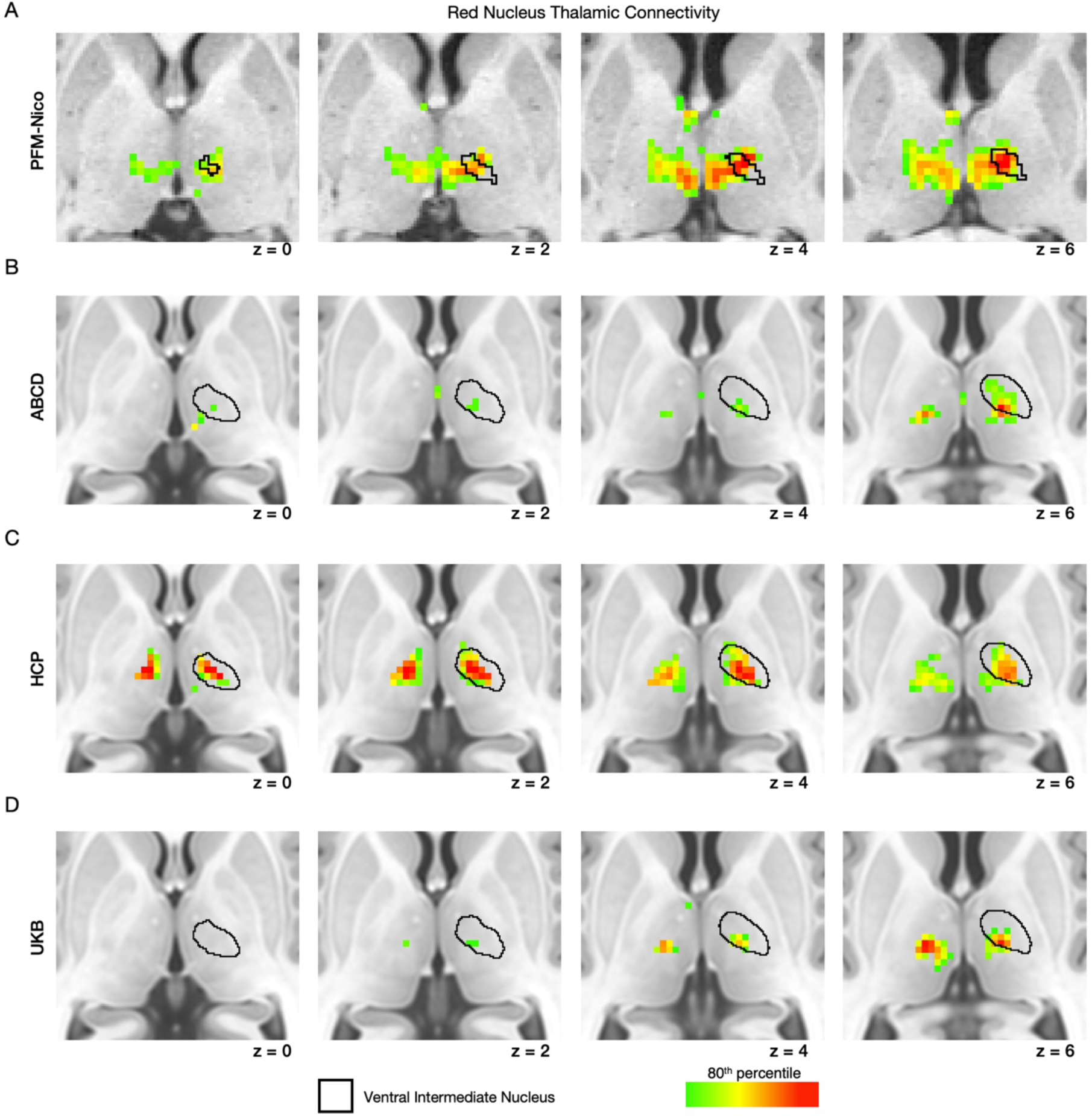
Thalamic connectivity of the red nucleus. Top 20% of red nucleus connections for the thalamus (MNI space) for PFM-Nico (A), ABCD study (n = 3,928) (B), HCP (812) (C), and UKB (n = 4,000) (D). Four different axial slices of the thalamus are shown (MNI space) overlaid on the subject’s structural image (A only). Thresholding is based on the top 20% of connections for the thalamus. The VIM (ventral intermediate) nucleus of the thalamus defined in an individual subject is shown in panel A. The nucleus outline based on a probabilistic map using THOMAS is shown in panels B-D. See supplemental figure 5 for additional participants.

### Red nucleus is not connected to effector-specific primary motor cortex

Evaluating the red nucleus as a whole, could potentially obscure a subregion of red nucleus with motor-effector connectivity. To further evaluate the hypothesis that red nucleus should have motor-effector network connectivity, we applied a winner-take-all approach to assign red nucleus voxels to networks based on cortical connectivity (Greene et al., 2020; Zheng et al., 2021). We found that almost no voxels had preferential motor-effector specific connectivity (foot, hand, mouth) (Fig 3, hollow triangles) in large group-averaged (Fig. 3, left) or individual-specific (Fig. 3, right) datasets. Most voxels were assigned to either CON, salience, or SCAN (Fig 3, filled circles).

**Figure 3:**
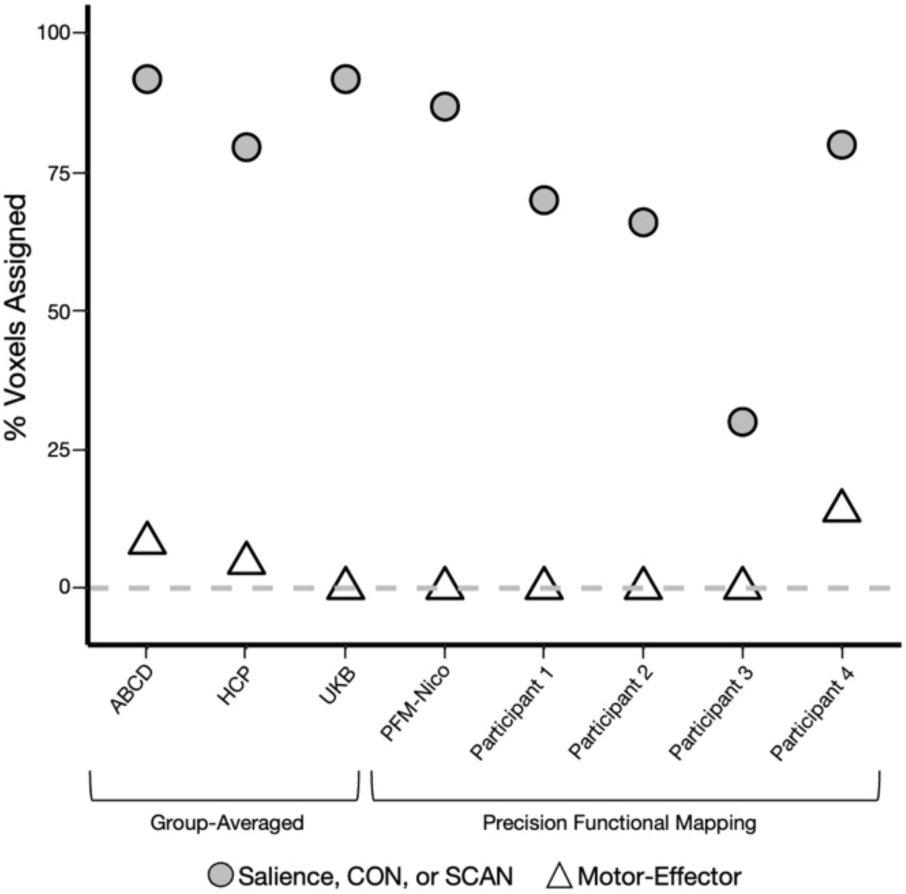
Action versus motor-effector network assignments of red nucleus voxels. Using winner-take-all red nucleus voxels were assigned to networks. The percentage of red nucleus voxels assigned to action related networks (salience network, CON, or SCAN) is shown with filled circles and the percent assigned to the three motor-effector networks is shown with hollow triangles. The left three columns show group-averaged data compared to the right five columns showing data from highly sampled individuals.

### Distinct ventral-lateral (salience) and dorsal-medial (cingulo-opercular) subdivisions

Winner-take-all assignments identified two sub-populations within the red nucleus, one with connectivity to the salience network and one to the CON/SCAN (SFig 7A). To delineate red nucleus subdivisions, we used agglomerative hierarchical clustering which grouped voxels based on functional connectivity with cortical networks (Gan et al., 2020; Greene et al., 2020). These analyses identified a dorsal-medial and ventral-lateral division of the red nucleus (Fig 4A, SFig 7:8, STable1). Comparing the functional connectivity of these two sub-divisions ((ventral-lateral [salience preference] - dorsal-medial [CON preference]); Zheng et al., 2021) demonstrated that the ventral-lateral division had stronger connectivity to the salience and parietal memory network (Gilmore et al., 2021; Zheng et al., 2021), while the dorsal-medial red nucleus had stronger connectivity to the CON and to SCAN regions within the precentral gyrus (Fig 4B,C, SFig 8A). Using preference for Salience or CON connectivity alone in a forced choice was able to identify the two red nucleus partitions (AUC>0.9). In support of this dorsal-medial/ventral-lateral partition of red nucleus, we also examined the correlation between red nucleus connectivity and specific cortical networks, revealing an obvious divide in network connectivity between CON and salience (Fig 4C:D, SFig 7B, SFig 8D). Neither subdivision displayed effector-specific motor connectivity, and as with connectivity of the entire red nucleus, dorsal-medial red nucleus precentral gyrus connectivity was strongest to SCAN regions (Fig 4B,C). Using the preference for salience or CON in group average datasets identified a similar ventral-lateral (salience) and dorsal-medial (CON/SCAN) division (SFig 7B, SFig 9).

**Figure 4:**
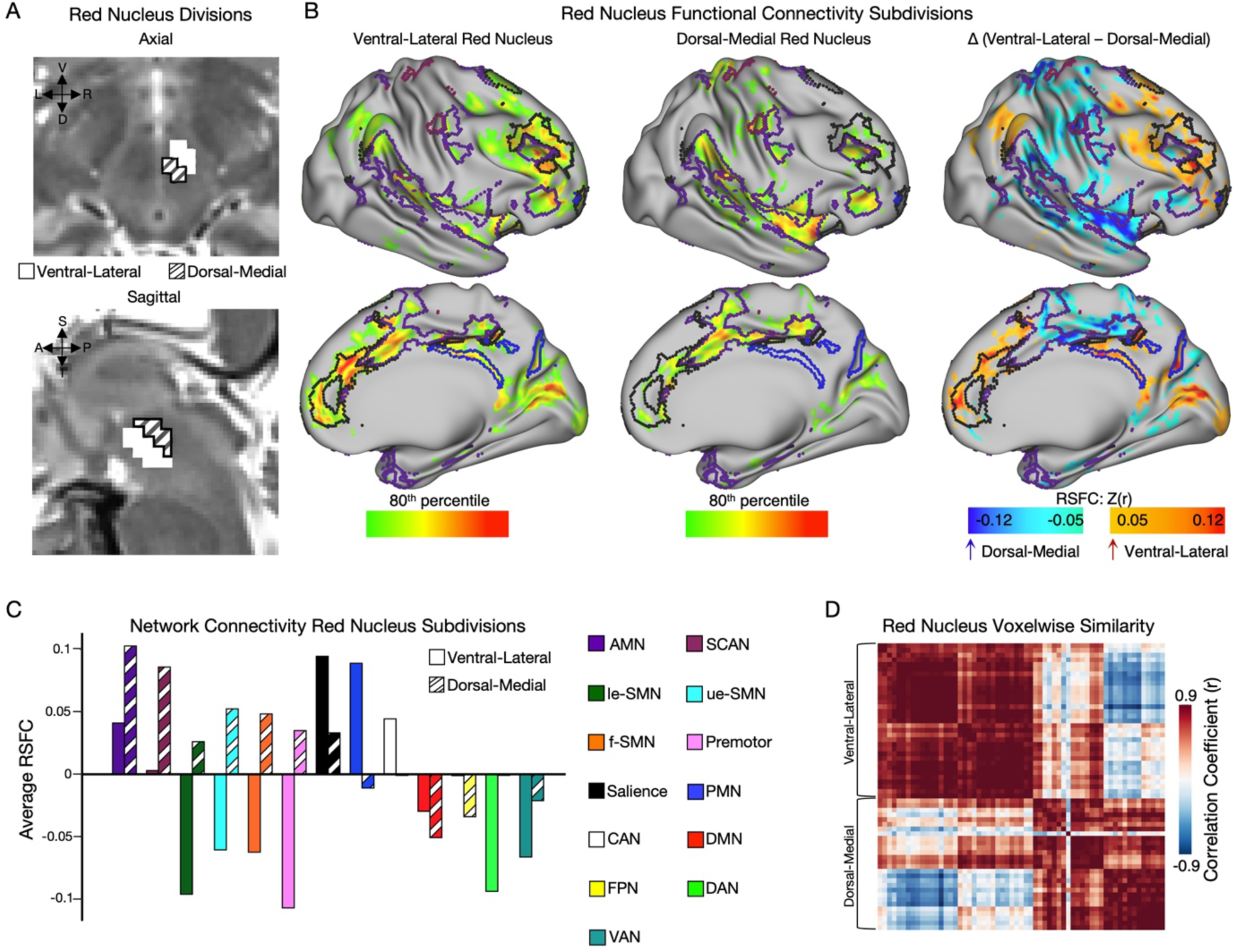
Functional connectivity subdivisions of the red nucleus. A) Anatomical display of dorsal-medial (hatched) and ventral-lateral (no fill) red nucleus subdivisions in exemplar participant (PFM-Nico) overlaid on T2w image. B) Strongest 20 percent of cortical RSFC for ventral-lateral (left) and dorsal-medial (middle) red nucleus subdivisions. Right most image shows the difference map between these two connectivity maps with greater ventral-lateral connectivity in red and greater dorsal-medial in blue. C) Average cortical RSFC organized by network for dorsal-medial (hatched) and ventral-lateral (no fill) subdivisions. D) Similarity (r) in network connectivity for each red nucleus voxel grouped into dorsal-medial and ventral-lateral divisions. For additional subjects/analyses see supplemental figures 6:8.

Given that there were two discrete subdivisions within the red nucleus favoring the salience network (ventral-lateral) or CON (dorsal-medial), we used each as separate seeds when determining subcortical connectivity. The ventral-lateral salience network favoring red nucleus partition was functionally connected to the VIM (SFig 10A). The ventral-lateral partition had peak cerebellar connectivity in lobule VI (SFig 10A). The dorsal-medial CON favoring partition was functionally connected to the mediodorsal nucleus of the thalamus (SFig 10B). This dorsal-medial red nucleus partition also had connectivity peaks in cerebellar lobule VIII, especially in VIIIb (SFig 10B).

## Discussion

### The human red nucleus is no longer a motor structure

The red nucleus is functionally connected to action-control (cingulo-opercular) and motivated behavior (salience) networks, but not to motor-effector networks. These results, combined with the reduction in the rubrospinal pathway in humans (Massion, 1988; Nathan & Smith, 1982), suggest that the red nucleus no longer controls movements in humans, arguing against the motor-control hypothesis. In fact, the red nucleus displayed no functional connectivity (or negative functional connectivity) with motor-effector networks, and motor cortex connectivity was restricted to SCAN regions (Gordon et al., 2023). The evolutionary principle of ‘exaptation’, where a trait serves a new function other than its original purpose, may be useful in understanding the red nucleus. The original function was coordinated extremity movement for locomotion, but the pyramidal system emerged and bipedalism made the rubrospinal pathway outdated for its prior functions (Ten Donkelaar, 1988). Instead of gradually disappearing, the once motor red nucleus was ultimately repurposed.

Multiple non-human primate studies have shown red nucleus structural connectivity to the motor cortex using tract tracing (Burman et al., 2000; H. G. Kuypers & Lawrence, 1967). Somewhat surprisingly, human red nucleus functional connectivity to effector-specific regions of motor cortex was small or negative. Instead, red nucleus functional connectivity to motor cortex appeared limited to the SCAN regions. This conflicts with motor-control models that position the red nucleus (and potentially the entire DRTT) as a system for the modification of motor-effectors in the cortex. Instead, the observed functional connectivity to SCAN is more consistent with the red nucleus’ original role in whole body coordination, an operation that activates SCAN (Gordon et al., 2023). One could predict that a re-analysis of non-human primate red nucleus tract tracing studies that factor in SCAN homologues in precentral gyrus would show red nucleus motor cortex connectivity to be specific to SCAN homologues, instead of motor-effector regions.

### Red nucleus is positioned to integrate reward and action

It remains an open question whether the salience and CON subdivisions of the red nucleus are strictly parallel or whether they might support the integration of reward/motivated-behaviors (hence salience network connectivity) and action-control (hence CON connectivity), allowing action plans to be regulated by motivation, even in the brainstem. This general framework is consistent with emerging perspectives that the brain must produce specific behavior in the context of motivated states (W. E. Allen et al., 2019). The red nucleus could serve as a tool to guide and rapidly adapt action execution based on changing salience information. In either scenario, it appears human action control includes repurposed motor nuclei, and highlights an evolutionary link between thinking/planning and movement (György Buzsáki, 2019; Llinás, 2002).

### Neural networks are an organizing principle in the brainstem

The realization that the red nucleus function seems to have shifted its role from quadrupedal locomotion to reward and action processes has broader implications. The brainstem has often been conceptualized as participating in two rigid hierarchies: a top-down control hierarchy where it passively receives and transmits top-down signals originating from the cortex (often motor commands); and a bottom-up sensory hierarchy where it passively receives and transmits sensory signals originating from the periphery. Instead, tract tracing indicates that neither of these perspectives are fully applicable to the red nucleus, given the small connection with the spinal cord. We speculate that the dominant representation of two cortical networks in the red nucleus indicates that neural networks are a principle of whole brain organization, and not just the cerebral cortex, in keeping with recent findings and perspectives (Chin et al., 2023; Gordon et al., 2020; Raut et al., 2021). How specific networks interact with the body, establishing the brain-body relationship (Chiel & Beer, 1997; Dum et al., 2019), is a topic that warrants further investigation and would likely have important implications to studies of affect and motivated behavior.

### Clinical targeting can benefit from red nucleus heterogeneity

Our improved understanding of human red nucleus connectivity and organization could reveal new targeting approaches for neuromodulation. Despite indications of red nucleus pathology in illnesses like Parkinson’s disease (Guan et al., 2017), we are aware of less than a half dozen studies that investigate red nucleus stimulation for disease, such as essential tremor or tremor-predominant Parkinson’s Disease. In one case, an electrophysiologic profile of the red nucleus was developed with the goal of avoiding the red nucleus in brain stimulation (Micieli et al., 2017). This is not without reason, as the side effect profile for red nucleus stimulation is alarming, because axons of the third cranial nerve pass through the red nucleus and red nucleus damage/stimulation can result in ocular disturbances (Lefranc et al., 2014; Leys et al., 1992). This proximity to oculomotor nuclei/axons may help to explain red nucleus functional connectivity with visual cortex (Fig 1B). Interestingly, the insertion of a macro-electrode into the red nucleus transiently reduced postural tremor in a single patient (Lefranc et al., 2014). Our findings suggest that using whole red nucleus connectivity (functional and potentially structural) could be a suboptimal approach for identifying thalamic targets. Instead, using the functional connectivity of each red nucleus division to the thalamus could prove to be an effective strategy for fine-tuning VIM or mediodorsal stimulation sites at an individual level for the treatment of tremor or pain (Meda et al., 2019) respectively.

### Salience and action control loops converge in the red nucleus

In conclusion, the absence of motor-effector functional connectivity and strong salience and cingulo-opercular network (CON) connectivity argues against the human red nucleus being a motor-control nucleus that indirectly influences motor-effector neurons in M1 to modify movement. The human red nucleus may form a node in a loop between the cortex and the cerebellum to integrate motivated behavior into action control, facilitating goal-directed behavior. The functional coupling of brainstem nuclei with higher-order action control networks indicates that neuroscience can benefit from taking a holistic approach to investigations of the brain (Chin et al., 2023).

## Methods

### Washington University participant for precision functional mapping (PFM-Nico)

The participant was 37 year old healthy adult male used in both the Midnight Scan Club ((Gordon et al., 2017); MSC02) and limb immobilization studies ((Newbold et al., 2020) ;SIC01), and the senior investigator of this current project (N.U.F.D.). This participant is referred to as precision functional mapping (PFM)-Nico.

### Cornell University participants for precision functional mapping

Four healthy adult participants (ages 29, 38, 24, and 31; all male) from a previously published study were used (Lynch et al., 2020). These participants are referred to as participant 1:4 in the manuscript. The previous study was approved by the Weill Cornell Medicine Instructional Review Board and each participant provided written informed consent. For additional details please see ref. (Lynch et al., 2020).

### UK Biobank (UKB)

We downloaded the group-averaged weighted eigenvectors from an initial group of 4,100 UKB participants aged 40-69 years (53% female) with 6-minute resting-state scans (https://www.fmrib.ox.ac.uk/ukbiobank/). Details of the acquisition and processing can be found at https://biobank.ctsu.ox.ac.uk/crystal/ukb/docs/brain_mri.pdf (Miller et al., 2016). This eigenvector file was mapped to the Conte69 surface template (Van Essen et al., 2012) using the ribbon-constrained method in Connectome Workbench (Glasser et al., 2013), following which the eigenvector time courses were cross-correlated.

### Adolescent Brain Cognitive Development (ABCD) Study

3,928 9-10-year-old participants (51% female), with at least 8 minutes of low-motion resting state data were used. In cases (e.g. Fig 1C) these subjects were split into two equal halves as described previously (Marek et al., 2022). Data processing was done with the ABCD-BIDS pipeline (NDA collection 3165; https://github.com/DCAN-Labs/abcd-hcp-pipeline). For additional details see: (Casey et al., 2018; Feczko et al., 2021; Marek et al., 2022).

### Human Connectome Project (HCP)

The group-averaged dense functional connectivity matrix for the HCP 1200 participants release, consisting of functional connectivity data for all 812 participants aged 22-35 years (410 female) with 60 minutes of resting-state fMRI, was downloaded from https://db.humanconnectome.org. For more information on the acquisition and processing see: (Glasser, Coalson, et al., 2016; Glasser et al., 2013; Glasser, Smith, et al., 2016; Smith et al., 2013).

### Preprocessing of PFM-Nico

PFM-Nico refers to a single participant (N.U.F.D) collected at Washington University in St. Louis. Imaging was performed using a Siemens TRIO 3T MRI scanner. Structural MRI was consisted of four T1-weighted images (sagittal acquisition, 224 slices, 0.8 mm isotropic resolution, [TE etc]) and four T2-weigthed images (sagittal [details]). Structural data were processed using previously described methods (Newbold et al., 2020) in which all T1w and T2w were separately averaged into a single structural file for functional processing and registration. Functional data acquisition was done using a multi-echo gradient-echo sequence consisting of nine 15-minute runs ([parameters]). In addition, 3 noise frames were acquired per run for noise reduction with distribution corrected (NORDIC) PCA, which was used to reduce thermal noise in functional data (Dowdle et al., 2023).

Optimal combination of multi-echo data and multi-echo independent component analysis (MEICA) denoising were performed using the tedana package version 0.0.11 (DuPre et al., 2021; Kundu et al., 2012, 2013). To promote reproducibility, we copy the automated methods description writeup as follows. TE-dependence analysis was performed on input data. An initial mask was generated from the first echo using nilearn’s compute_epi_mask function. An adaptive mask was then generated, in which each voxel’s value reflects the number of echoes with ’good’ data. A two-stage masking procedure was applied, in which a liberal mask (including voxels with good data in at least the first echo) was used for optimal combination, T2*/S0 estimation, and denoising, while a more conservative mask (restricted to voxels with good data in at least the first three echoes) was used for the component classification procedure. A monoexponential model was fit to the data at each voxel using log-linear regression in order to estimate T2* and S0 maps. For each voxel, the value from the adaptive mask was used to determine which echoes would be used to estimate T2* and S0. Multi-echo data were then optimally combined using the T2* combination method (Posse et al., 1999).

Principal component analysis based on the PCA component estimation with a Moving Average(stationary Gaussian) process (Li et al., 2007) was applied to the optimally combined data for dimensionality reduction. The following metrics were calculated: kappa, rho, countnoise, countsigFT2, countsigFS0, dice_FT2, dice_FS0, signal-noise_t, variance explained, normalized variance explained, d_table_score. Kappa (kappa) and Rho (rho) were calculated as measures of TE-dependence and TE-independence, respectively. A t-test was performed between the distributions of T2*-model F-statistics associated with clusters (i.e., signal) and non-cluster voxels (i.e., noise) to generate a t-statistic (metric signal-noise_z) and p-value (metric signal-noise_p) measuring relative association of the component to signal over noise. The number of significant voxels not from clusters was calculated for each component. Independent component analysis was then used to decompose the dimensionally reduced dataset. The following metrics were calculated: kappa, rho, countnoise, countsigFT2, countsigFS0, dice_FT2, dice_FS0, signal-noise_t, variance explained, normalized variance explained, d_table_score. Kappa (kappa) and Rho (rho) were calculated as measures of TE-dependence and TE-independence, respectively. A t-test was performed between the distributions of T2*-model F-statistics associated with clusters (i.e., signal) and non-cluster voxels (i.e., noise) to generate a t-statistic (metric signal-noise_z) and p-value (metric signal-noise_p) measuring relative association of the component to signal over noise. The number of significant voxels not from clusters was calculated for each component. Next, component selection was performed to identify BOLD (TE-dependent), non-BOLD (TE-independent), and uncertain (low-variance) components using the Kundu decision tree (v2.5 (Kundu et al., 2013)). This workflow used numpy (Van Der Walt et al., 2011), scipy (Jones et al., 2001), pandas (McKinney, 2010), scikit-learn (Pedregosa et al., 2011), nilearn, and nibabel (Brett et al., 2019). This workflow also used the Dice similarity index (Dice, 1945; Sorensen, 1948).

For every run of BOLD data, we manually inspected the noise/signal classification from MEICA and adjusted classification where needed. This strategy of manual inspection is viable in the context of small sample studies like ours, and is a major strength of the PFM approach. Only components classified as signal were used for all analyses. Based on the 6 rigid body parameters derived via retrospective motion correction, we calculated frame-wise displacement (FD (Power et al., 2012)). Motion parameters were low-pass filtered (threshold set at 0.1 Hz) before FD computation so as to reduce the impact of respiratory artifact of estimates of head motion (Fair et al., 2020). To identify high motion frames, we set a threshold of 0.08 mm on the FD vector. Global signal was calculated as the average of all voxels within a brain mask. Following optimal combination and MEICA, data underwent temporal bandpass filtering with frequencies between 0.005 Hz and 0.1 Hz being retained. Global signal and its first derivative constituted the only nuisance regressors. Following noise correction, cortical data were projected onto a surface using a previously described approach (Gordon et al., 2017). Data were smoothed with a geodesic 2D (surface) or Euclidean 3D (volumetric) Gaussian kernel of *σ* = 2.55 mm. Volumetric smoothing was done within each subvolume including bilateral red nuclei (described in *manual tracing of the red nucleus*).

### Improving brainstem signal-to-noise ratio

The brainstem is the most difficult part of the brain to acquire high quality functional neuroimaging data. Distance from the head coils inherently makes the SNR lower here than in the cortex. Additionally, the optimal echo time is different in the brainstem (and varies across the brainstem) than the cerebral cortex, in part due to high concentration of iron. Given that most studies optimize scanning parameters for the cortex, common scanning parameters are poorly suited for the brainstem. Also, we encountered sources of noise at the individual level that were difficult to characterize with standard denoising with motion and anatomical regressors. In total, these limitations with current brainstem imaging required a specialized denoising strategy that would allow us to acquire high quality cortical data along with brainstem data. The first part of this strategy was the implementation of a recently developed thermal denoising approach called NORDIC (Dowdle et al., 2023), which greatly reduces noise. By acquiring multi-echo data and employing optimal combination of echoes on a voxelwise manner, we were able to have an optimized echo time for both cortical and brainstem data. Also, MEICA allows for a substantial improvement in SNR (Lynch et al., 2020). We utilized MEICA and manually modified noise components on a run-by-run level, in a process that would be excessively burdensome for large sample size studies, but is viable in a PFM framework. Finally, we collected a far greater amount of data on an individual level than the vast majority of groups, allowing for a ‘brute force’ approach for SNR improvement. When we were incapable of applying these denoising strategies (in the case of group averaged datasets and single echo datasets) we relied on massive sample sizes to improve SNR. In total, our results allowed us to overcome the most pressing issue on brainstem functional imaging, namely, the low SNR. The strategies employed in the paper demonstrate the feasibility of brainstem neuroimaging and can be used to investigate clinically relevant structures like the substantia nigra and periaqueductal grey.

### Defining the red nucleus

Unlike many brainstem nuclei, the red nucleus is clearly visible on T2-weighted images as a hypo-intensity (Fig 1A). Individual level red nucleus was hand drawn on T2-weighted native space images (Fig 1A) by a single experimenter (S.R.K.) and transformed to MNI space for subsequent analyses. Publicly available brainstem atlases were used as a reference for the red nucleus to assist in manual drawing (brainstem navigator atlas https://www.nitrc.org/projects/brainstemnavig (Bianciardi et al., 2015)). For group average datasets, the red nucleus was again hand drawn but on a high resolution T2-weighted MNI template image.

### Cortical network identification

The Infomap algorithm (Rosvall & Bergstrom, 2008 : https://www.mapequation.org/) was used to assign vertices to communities, and the resulting communities were then assigned a network identity based on similarity to known group-average networks. The consensus network assignment, computed by aggregating across thresholds, was used as the cortical resting state networks (see SFig 2 for example assignments). The original 17 networks set described in MSC (ref to msc paper) was recently amended to account for the SCAN (Gordon et al., 2023).

### Red nucleus functional connectivity

Using the red nucleus as a seed we averaged the timeseries of red nucleus voxels to create a red nucleus timeseries and correlated this to brain. In cases of HCP and UKB, we instead averaged over rows in the dense connectivity file corresponding to the red nucleus. When determining functional connectivity to red nucleus subdivisions, we simply repeated this process, but for the subdivision instead of the whole red nucleus.

### Winner-take-all analysis of red nucleus voxels

We used a previously established approach for assigning red nucleus voxels to bilateral cortical networks (Zheng et al., 2021). Described briefly, a voxel was assigned to the network that it had the largest correlation to, so long as that correlation was greater than zero. We excluded three sensory networks, two visual and one auditory, from possible assignment, because the red nucleus is not believed to be involved in these processes, and because potential assignment to these three networks would be likely artifactual potentially owing to partial volume effects with the third cranial nerve which passes through the red nucleus. Additionally, inconsistent and small Infomap cortical assignment to the anterior and posterior medial temporal networks led us to exclude these two networks as well. In total, there were 13 networks that red nucleus voxels could be assigned.

### Clustering

Clustering of the red nucleus was based on cortical connectivity, specifically the correlation between each red nucleus voxel and the 13 bilateral resting state networks similar to previous clustering approaches to other subcortical structures (Greene et al., 2020). We used hierarchical clustering on the Euclidean distance between cortical connectivity strength with Ward’s method (Murtagh & Legendre, 2014; Ward Jr, 1963). Using the NBclust R package we assessed clustering performance with the number of clusters ranging from 2 to 13 using more than 20 metrics (Charrad et al., 2014). For each number of clusters, a score for all clustering metrics was computed, and cluster performance was ranked (e.g. the number of clusters with the largest silhouette index scored a rank of 1). For each metric, a number of clusters “won” when it had the best performance for that specific clustering metric. The number of clusters chosen was based on a majority rule where the number of clusters with the most total victories (first place for each metric) was determined to be the best overall.

### Thalamus segmentation

The Thalamus-Optimized Multi-Atlas Segmentation (THOMAS v 2.1) (Jh et al., 2019) is a method for identification of nuclei, particularly the ventral intermediate nucleus that has been colocalized with the segment labelled the ventral part of the Ventro-Lateral-Posterior nucleus (Su et al., 2020).

To segment the thalamic nuclei on our precision mapping participant, we used the hips_thomas.csh function from the version 2. 1 that has been validated for use of T1 acquisition only (Pfefferbaum et al., 2023) and that is available on docker (https://github.com/thalamicseg/hipsthomasdocker). We used the average T1 acquisition that has been produced for the registration of all functional data. For group averaged data we used an MNI space transformation of a probabilistic THOMAS segmentation (https://zenodo.org/record/5499504).

### Projecting to the cerebellum

Cerebellar connectivity values were mapped onto a cerebellar flat map with the SUIT toolbox (Diedrichsen & Zotow, 2015).

### Statistics

We used a rotation-based null model to test if red nucleus connectivity was selective for networks or random (e.g. Fig 2B). In this approach, cortical resting state networks were rotated by a random amount around a spherical expansion of the cortical surface 1000 times (Gordon et al., 2016). For each rotation, we calculated the measure of interest (e.g. red nucleus connectivity strength to the rotated network). A p-value was calculated by comparing the true value against the values obtained through random rotations. When multiple comparisons across networks was performed, a Bonferroni correction for network number was used to control the false positive rate.

### Visualization

The distribution of functional connectivity values differs based on dataset, in part owing to different denoising decisions. Therefore, direct numerical comparisons across datasets is not appropriate. Thus, to facilitate comparison, in almost all figures, RSFC values were thresholded to be the top 20 percent of the given image. Supplemental Figure 2 demonstrates that this threshold does not obscure red nucleus connectivity. In group averaged subcortical data we noticed a pattern in which the edges of structures (e.g. thalamus) were the most likely to contain extreme values. Even an inspection of dense connectivity matrices shows an obvious effect of extreme values around the edge of volume structures. It is not entirely clear why this is the case. One possibility is that sub-volume constrained averaging may cause edge voxels to be noisier because they are averaged with fewer voxels, thus promoting extreme values. Thus, we excluded edge voxels from the thalamus exclusively for group average datasets. We accomplished this by minimally eroding the thalamus ROI by 3 mm. When examining Cornell data in the thalamus, we noticed that red nucleus signal was being smoothed into the ventral thalamus, leading to extremely large and erroneous connectivity values. To address this, we masked out thalamus voxels that were included in a 5 mm dilation of the bilateral red nucleus.

## Acknowledgements

This work was supported by NIH grants T32DA007261 (S.R.K), MH120989 (C.J.L.), MH120194 (J.T.W.), NS123345 (B.P.K.), NS098482 (B.P.K.), NS124789 (S.A.N.), MH120194 (J.T.W.), DA047851 (C.J.L.), MH118388 (C.J.L.), MH114976 (C.J.L.), MH129616 (T.O.L.), DA041148 (D.A.F.), DA04112 (D.A.F.), MH115357 (D.A.F.), MH096773 (D.A.F. and N.U.F.D.), MH122066 (E.M.G., D.A.F. and N.U.F.D.), MH121276 (E.M.G., D.A.F. and N.U.F.D.), MH124567 (E.M.G., D.A.F. and N.U.F.D.), NS129521 (E.M.G., D.A.F. and N.U.F.D.), and NS088590 (N.U.F.D.); by Center for Brain Research in Mood Disorders; by Eagles Autism Challenge; by the Dystonia Medical Research Foundation (S.A.N.); by the National Spasmodic Dysphonia Association (E.M.G. and S.A.N.); by the Taylor Family Foundation (T.O.L.); by the Intellectual and Developmental Disabilities Research Center (N.U.F.D.); by the Kiwanis Foundation (N.U.F.D.); by the Washington University Hope Center for Neurological Disorders (E.M.G., B.P.K. and N.U.F.D.); and by Mallinckrodt Institute of Radiology pilot funding (E.M.G. and N.U.F.D.).

