## Supplemental Figures for "The brainstem’s red nucleus was evolutionarily upgraded to support goal-directed action"

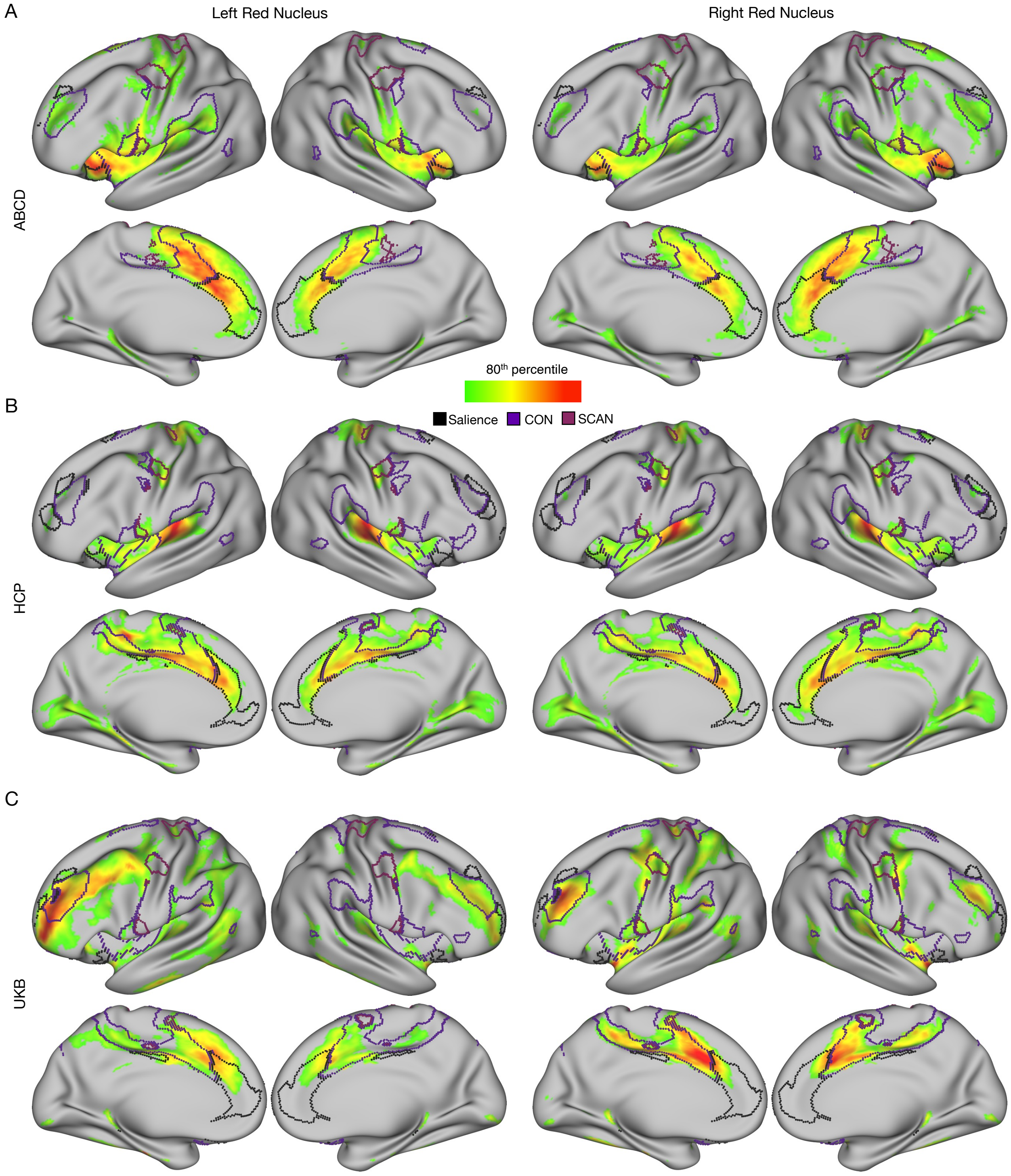


**Supplemental 1: Large dataset group averaged red nucleus cortical functional connectivity.** Red nucleus connectivity for ABCD study (n = 3,928) (A), HCP (n = 812) (B), and UKB (n = 4,000) (C) for left (left side of figure) and right (right side of figure) red nucleus. Network boundaries for salience (black), SCAN (mauve), and CON (purple) specific to each dataset are overlaid. CON (cingulo-opercular network; purple); SCAN (somato-cognitive action network; mauve). Thresholding is based on the top 20% of connections for the cortex.

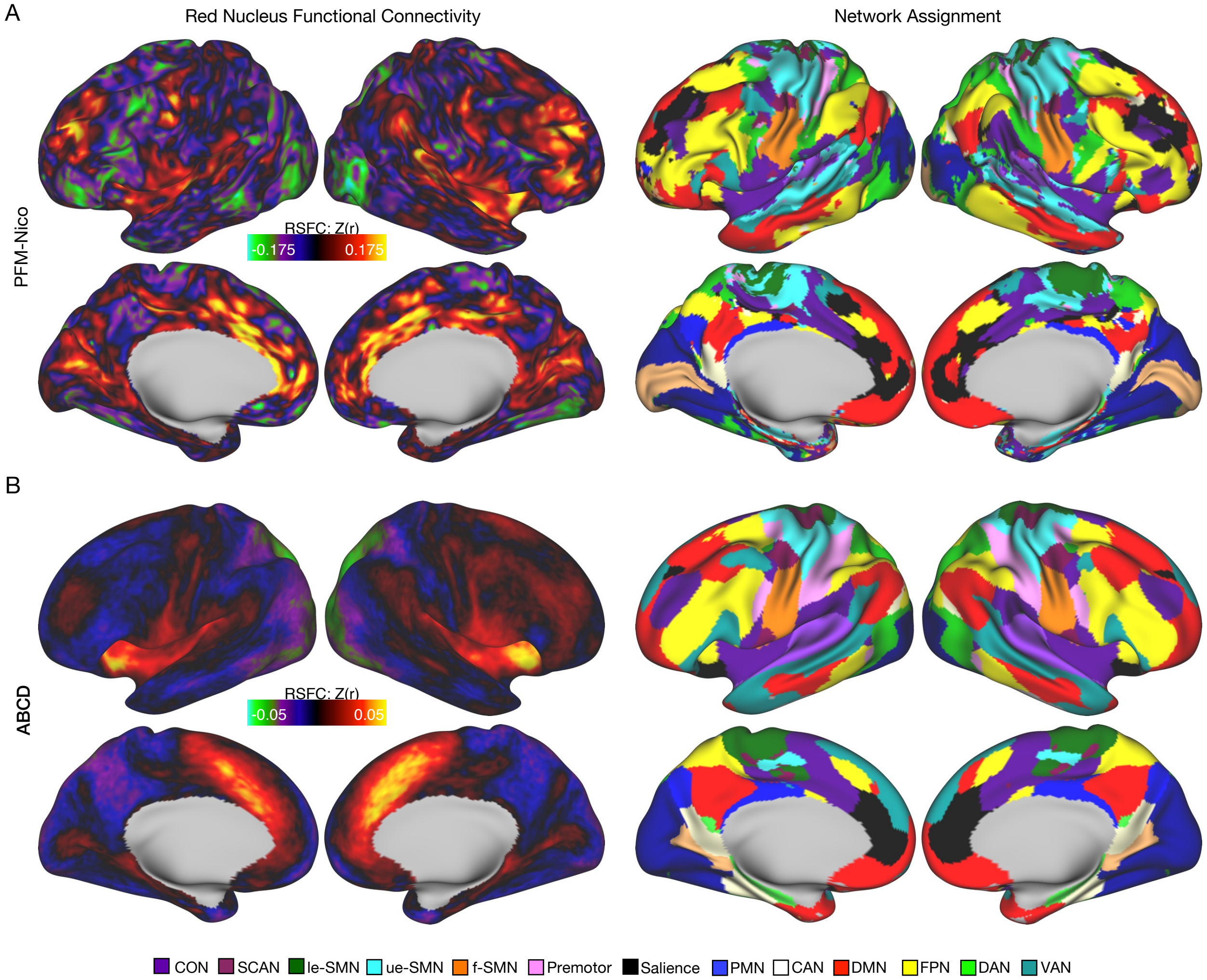


**Supplemental 2: Unthresholded red nucleus cortical functional connectivity.** Unthresholded right red nucleus functional connectivity for PFM-Nico (A, left) with corresponding individual specific network assignment (A, right). Panel B indicates the same as in panel A, but for the ABCD study. CON (cingulo-opercular network; purple); SCAN (somato-cognitive action network; mauve), le-SMN (lower-extremity somatomotor network; forest green), ue-SMN (upper-extremity somatomotor network; cyan), f-SMN (face somatomotor network; orange), PMN (posterior memory network; blue), CAN (contextual association network; white), DMN (default mode network; red), FPN (fronto-parietal network; yellow), DAN (dorsal attention network; neon green); VAN (ventral attention network; teal).

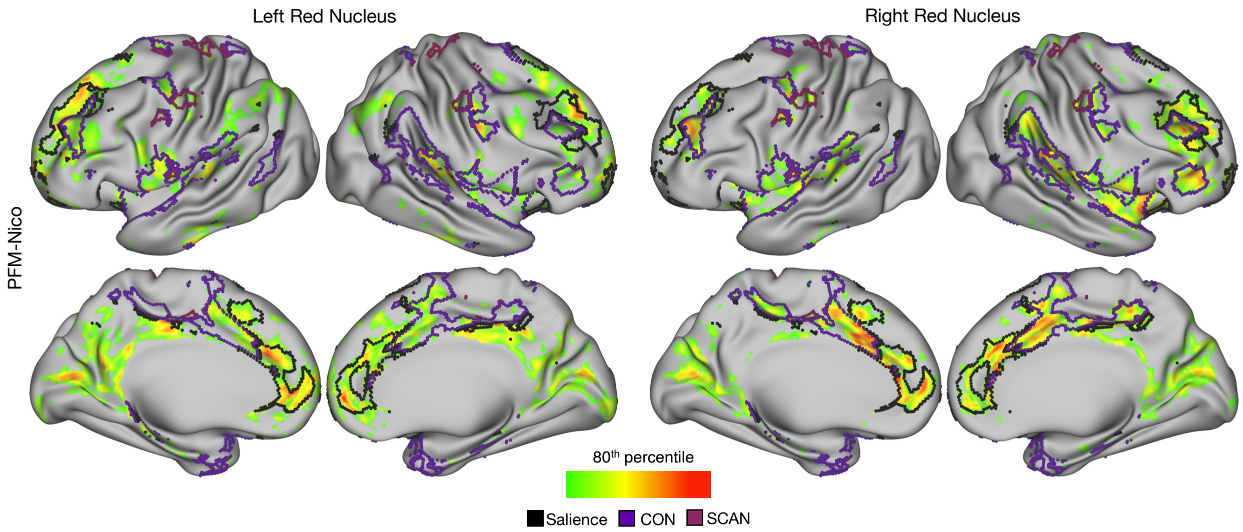


**Supplemental 3: Red nucleus resting state functional connectivity laterality.** Red nucleus connectivity for PFM-Nico for the left RN (shown on left) and the right RN (shown on right). CON (cingulo-opercular network; purple); SCAN (somato-cognitive action network; mauve). Thresholding is based on the top 20% of connections for the cortex.

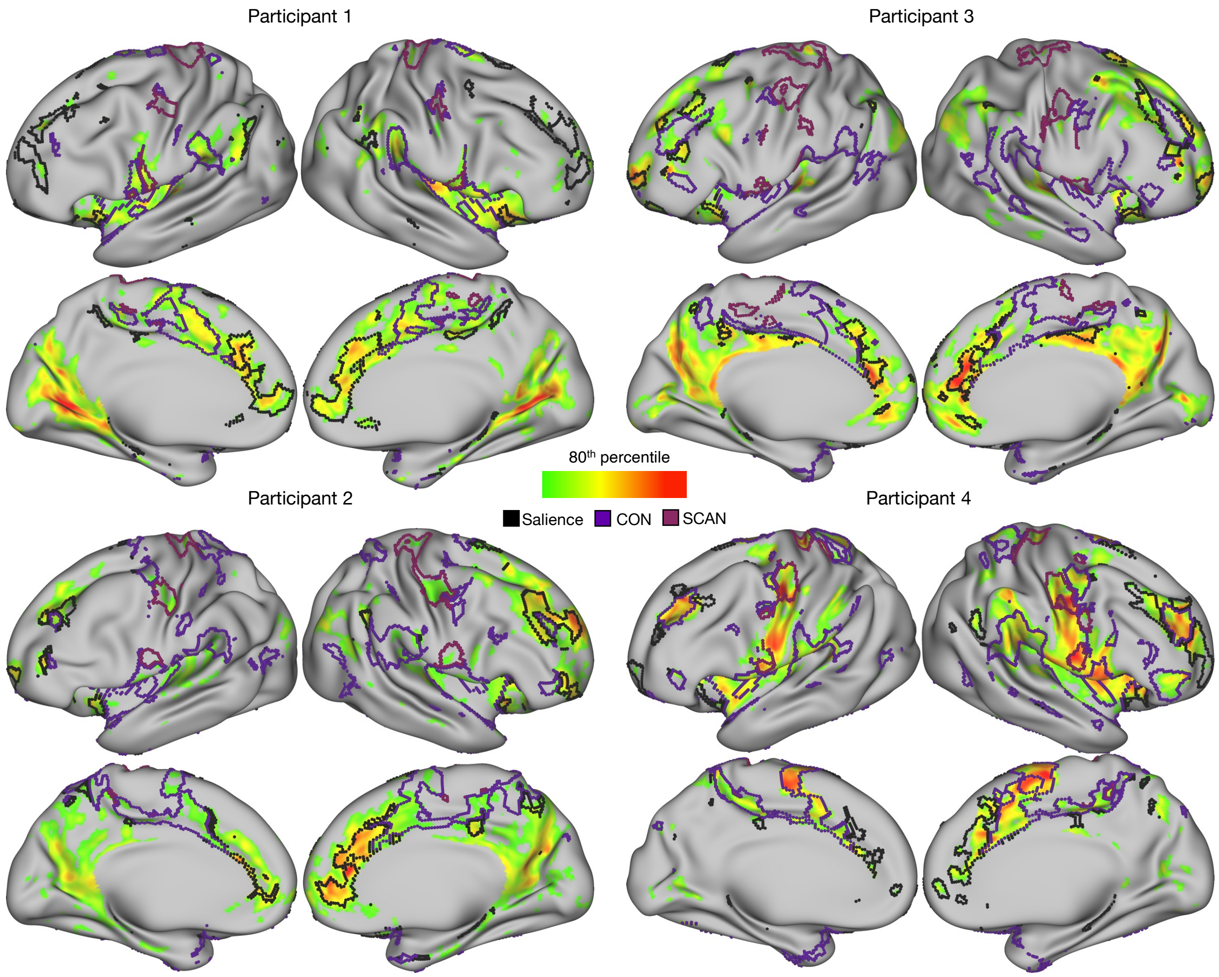


**Supplemental 4: Individual-specific precision functional mapping of red nucleus resting state functional connectivity – Cornell data.** Red nucleus connectivity for right side for four additional highly sampled subjected with multi-echo independent component analysis (MEICA) denoising. CON (cingulo-opercular network; purple); SCAN (somato-cognitive action network; mauve). Thresholding is based on the top 20% of connections for the cortex.

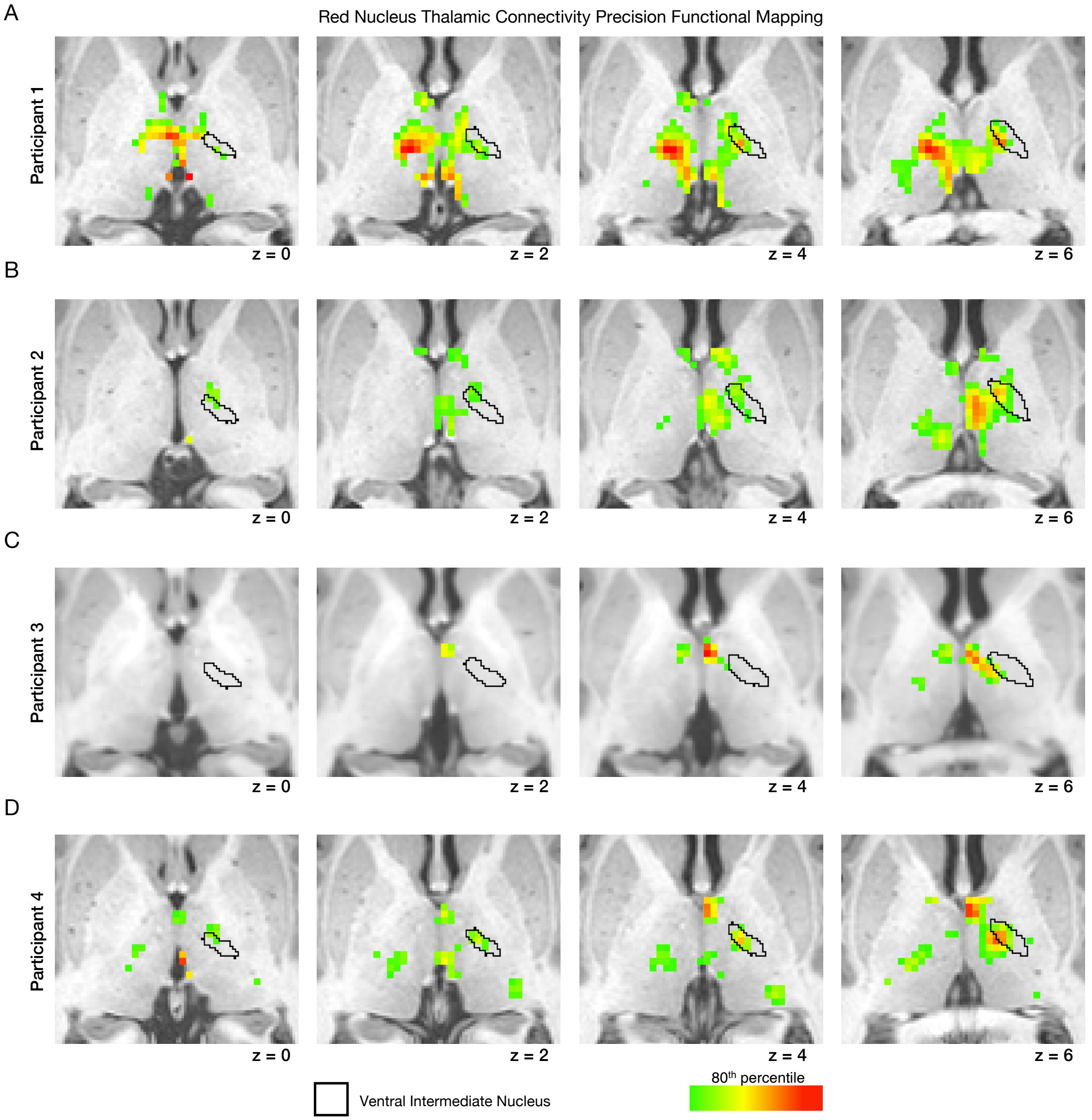


**Supplemental 5: Thalamic connectivity of the red nucleus in highly sampled individuals – Cornell data.** Top 20% of RN connections for the thalamus (MNI space) for four additional highly sampled subjects (A-D). Four different axial slices of the thalamus are shown (MNI space) overlaid on the subject's structural image. Thresholding is based on the top 20% of connections for the specific thalamus. The ventral intermediate nucleus was identified in an individual specific manner.

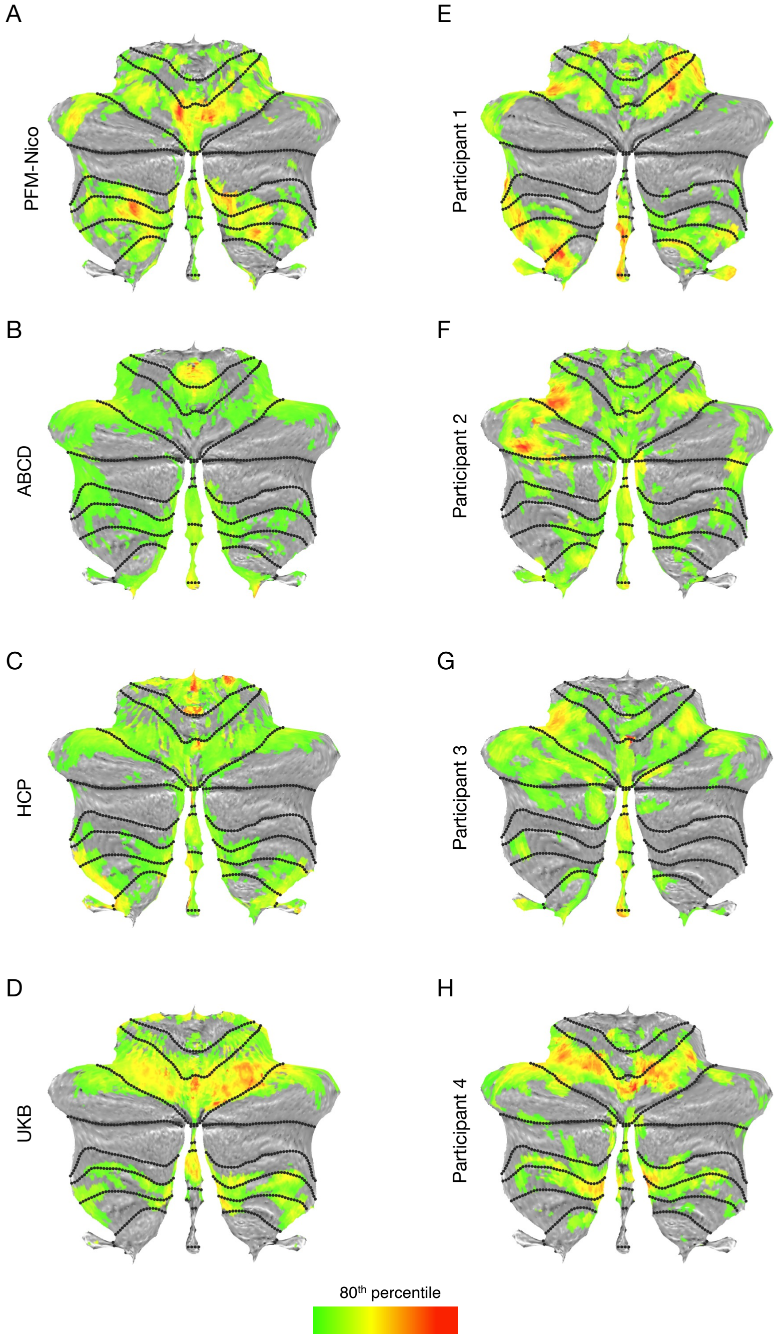


**Supplemental 6: Red nucleus cerebellar connectivity.** Cerebellar connectivity for the RN shown on a cerebellar flat map for PFM-Nico (A), ABCD (B, n=3,928), HCP (C, n=812), and UKB (D, n=4,000), Participants 1:4 (E:H). Thresholding is based on the top 20% of connections for the cerebellum.

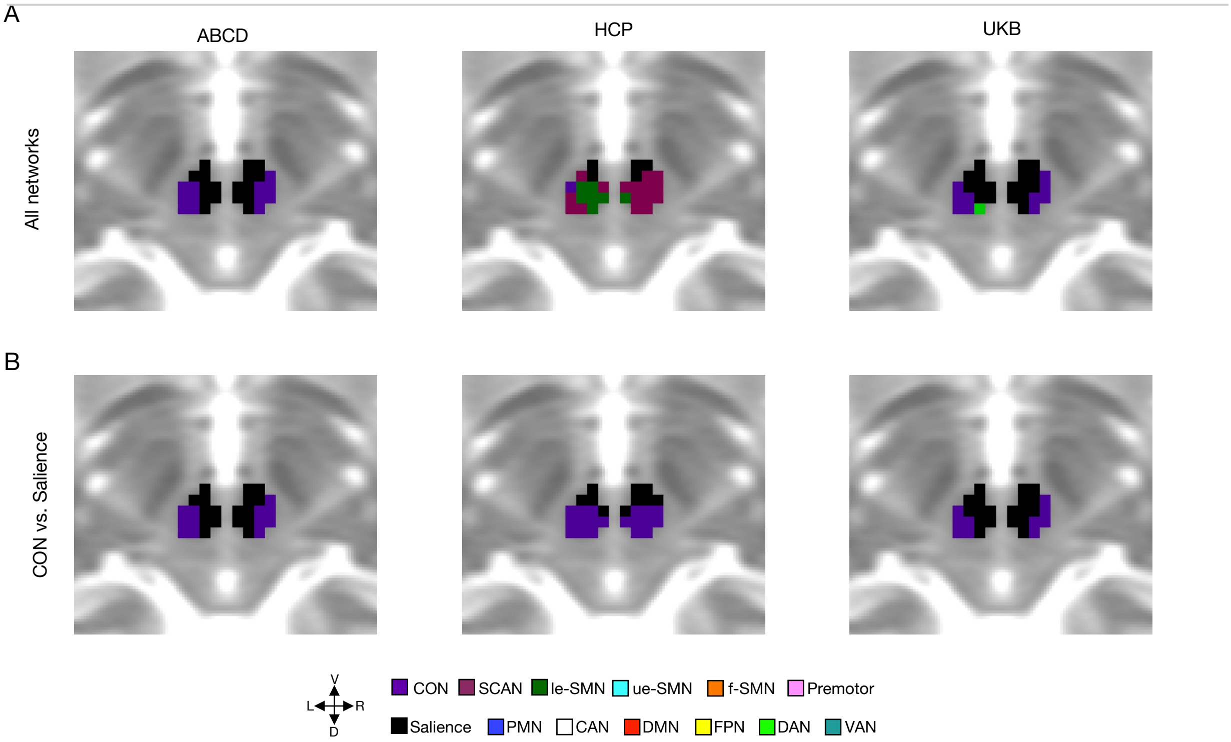


**Supplemental 7: Winner-take-all (WTA) red nucleus network assignments for group-averaged data.** WTA was used to assign red nucleus voxels in group average datasets to cortical networks. A) WTA assignment for all networks for ABCD (left), HCP (middle), and UKB (right). The force choice was restricted to CON or salience in panel B for the same three datasets. Shown at MNI space axial slice -8. CON (cingulo-opercular network; purple); SCAN (somato-cognitive action network; mauve), le-SMN (lower-extremity somatomotor network; forest green), ue-SMN (upper-extremity somatomotor network; cyan), f-SMN (face somatomotor network; orange), PMN (posterior memory network; blue), CAN (contextual association network; white), DMN (default mode network; red), FPN (fronto-parietal network; yellow), DAN (dorsal attention network; neon green); VAN (ventral attention network; teal).

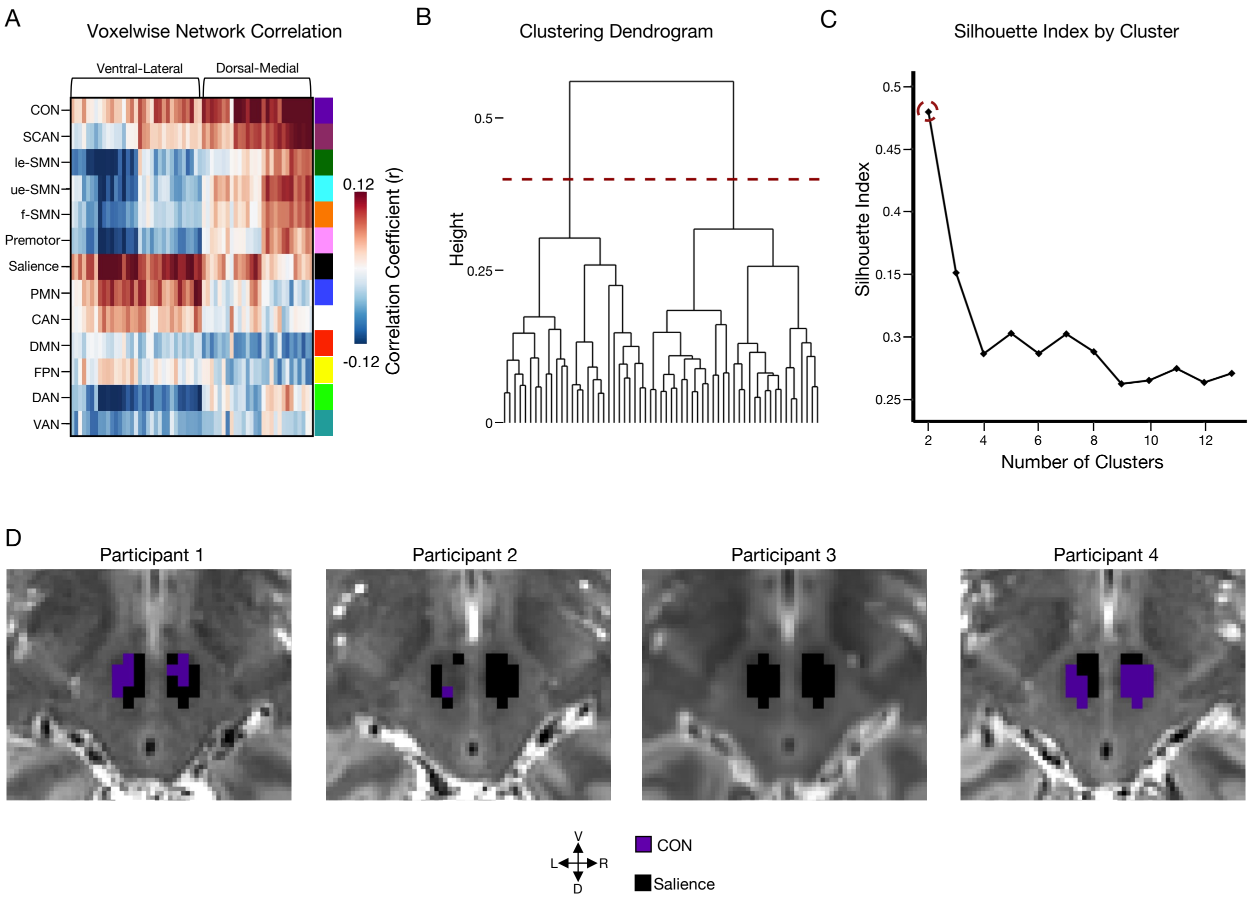


**Supplemental 8: Clustering decision support.** A) The correlation between each red nucleus voxel and the cortical networks organized into ventral-lateral and dorsal-medial divisions. B) The dendrogram resulting from hierarchical clustering. C) The silhouette index over different number of clusters highlighting two clusters was the best solution. Additional clustering metrics shown in STable1. D) Winner-take-all forced choice between salience and CON for additional participants shown at MNI axial slice -8. Lack of assignment for participant two indicates that no voxel reached threshold requirement for either network. CON (cingulo-opercular network; purple); SCAN (somato-cognitive action network; mauve), le-SMN (lower-extremity somatomotor network; forest green), ue-SMN (upper-extremity somatomotor network; cyan), f-SMN (face somatomotor network; orange), PMN (posterior memory network; blue), CAN (contextual association network; white), DMN (default mode network; red), FPN (fronto-parietal network; yellow), DAN (dorsal attention network; neon green); VAN (ventral attention network; teal).

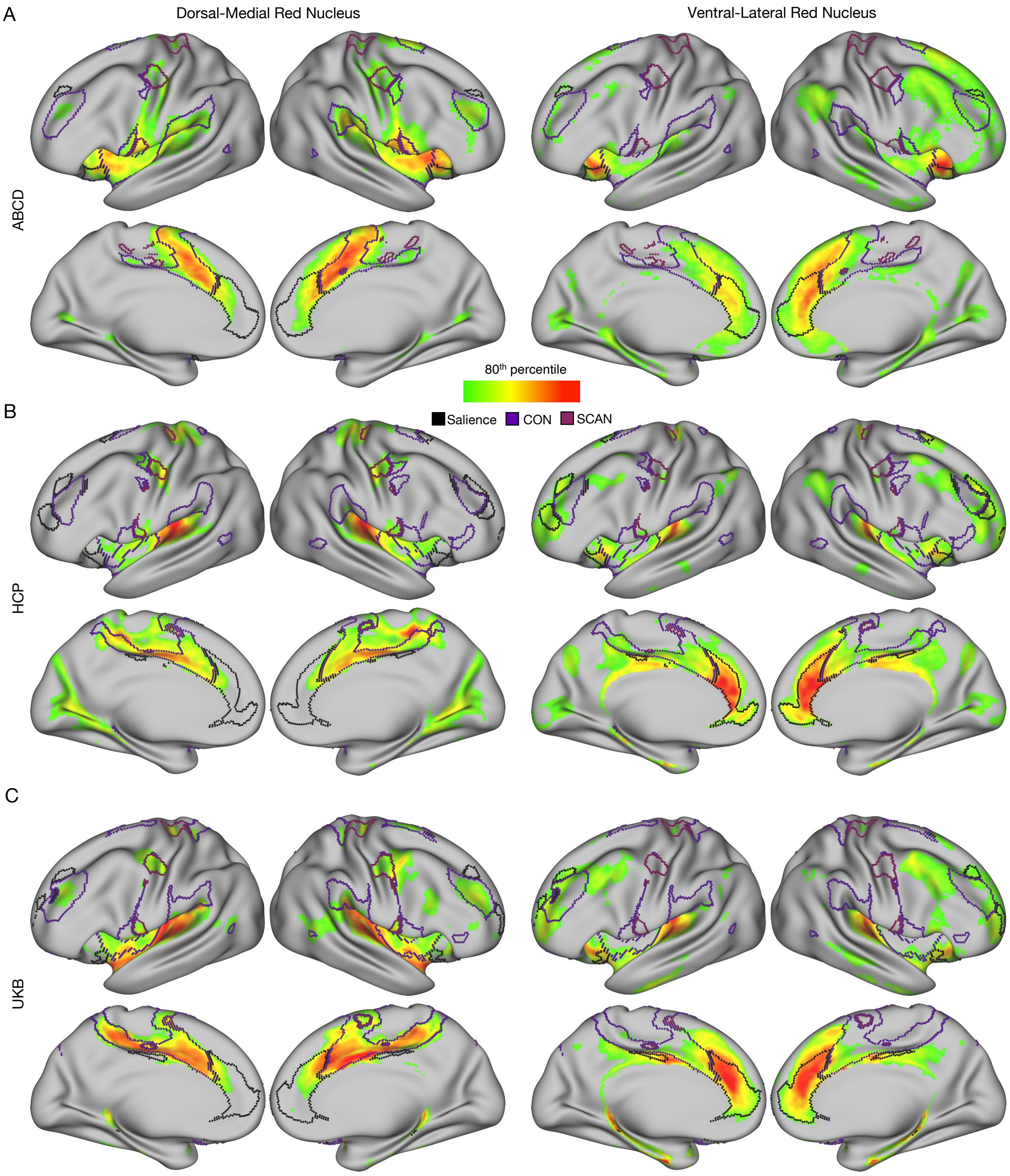


**Supplemental 9: Red nucleus subdivisions based on resting state functional connectivity in group-averaged data.** The dorsal-medial and ventral-lateral red nucleus were identified based on a forced choice between salience network (caudal) and CON (rostral) in the three group datasets. The left side shows connectivity for the dorsal-medial division for ABCD (A, n=3928), HCP (B, n=812), and UKB (C, n=4,000). The right side of the figure shows ventral-lateral divisions for these three datasets. CON (cingulo-opercular network; purple); SCAN (somato-cognitive action network; mauve). Thresholding is based on the top 20% of connections for the cortex.

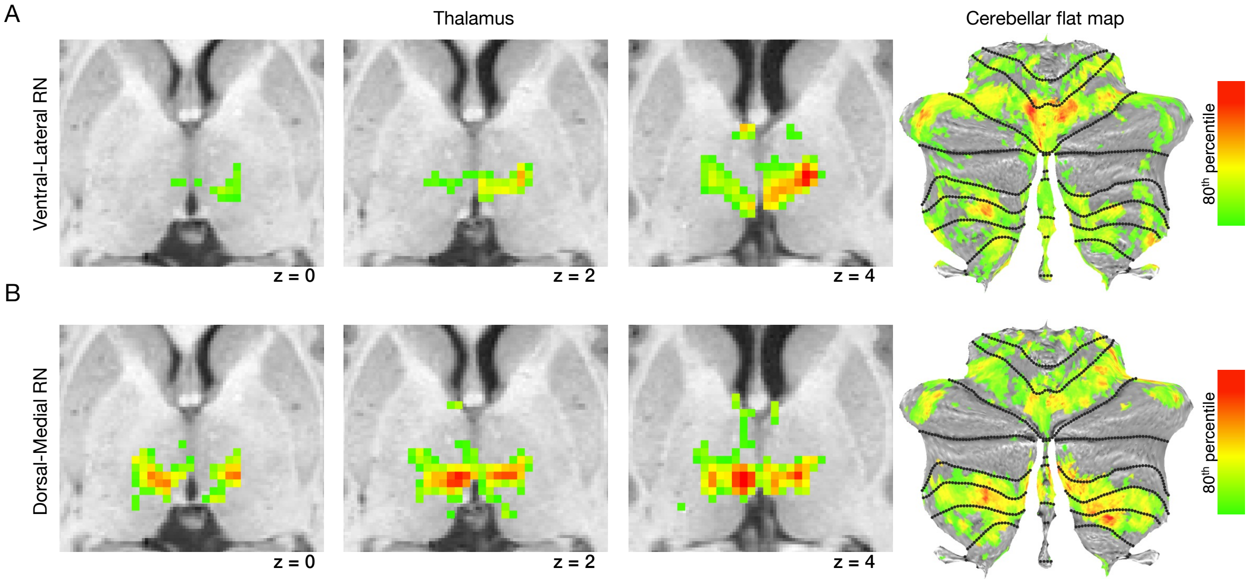


**Supplemental 10: Subcortical and cerebellar connectivity of red nucleus subdivisions in PFM-Nico.** Dorsal-medial (A) and ventral-lateral (B) red nucleus connectivity to the thalamus and cerebellum. Three axial slices of the thalamus are shown (MNI space) overlaid on the participant's T1w MRI. Thresholding is based on the top 20% of connections for the specific structure (thalamus or cerebellum).
