## Supplemental Table for "The brainstem’s red nucleus was evolutionarily upgraded to support goal-directed action"

| | 2 | 3 | 4 | 5 | 6 | 7 | 8 | 9 | 10 | 11 | 12 | 13 |
|---|---|---|---|---|---|---|---|---|---|---|---|---|
| KL | 8.852 | 0.8754 | 2.0771 | 1.6153 | 1.3088 | 1.0001 | 1.2884 | 1.1614 | 1.1771 | 1.0819 | 1.0686 | 0.9998 |
| CH | 92.554 | 64.0455 | 60.4721 | 54.7912 | 49.6614 | 45.6332 | 43.1456 | 40.812 | 38.8249 | 37.0271 | 35.5238 | 34.2604 |
| Hartigan | 14.6943 | 17.6122 | 9.9605 | 6.8497 | 5.5421 | 5.5875 | 4.5914 | 4.0913 | 3.5984 | 3.3865 | 3.2234 | 3.2541 |
| CCC | 12.6296 | 8.6493 | 6.9637 | 5.9991 | 4.8854 | 3.7746 | 4.0294 | 4.1139 | 4.1386 | 4.0889 | 4.0311 | 3.9699 |
| Scott | 306.8029 | 355.7281 | 418.0063 | 490.6359 | 532.2683 | 604.4944 | 720.2539 | 802.7899 | 824.4586 | 878.8139 | 922.9221 | 957.8301 |
| TrCovW | 0.0121 | 0.0063 | 0.0032 | 0.0022 | 0.0017 | 0.0014 | 0.0012 | 0.0011 | 0.0009 | 0.0007 | 0.0006 | 0.0006 |
| TraceW | 0.7387 | 0.5914 | 0.4536 | 0.3862 | 0.3441 | 0.3126 | 0.2833 | 0.2607 | 0.2417 | 0.2257 | 0.2114 | 0.1984 |
| Friedman | 637.2076 | 706.5274 | 750.0513 | 815.0031 | 844.1699 | 975.809 | 1137.733 | 1218.23 | 1237.345 | 1294.445 | 1378.604 | 1398.358 |
| Rubin | 4.4179 | 5.5182 | 7.1939 | 8.451 | 9.4847 | 10.4404 | 11.5207 | 12.5187 | 13.5037 | 14.4565 | 15.4356 | 16.451 |
| Cindex | 0.3917 | 0.3562 | 0.3484 | 0.3333 | 0.3637 | 0.0683 | 0.062 | 0.0569 | 0.0552 | 0.0535 | 0.0525 | 0.0507 |
| DB | 0.7973 | 1.1105 | 1.1453 | 1.154 | 1.2277 | 1.049 | 1.1441 | 1.2508 | 1.1837 | 1.1115 | 1.1078 | 1.079 |
| Silhouette | 0.4796 | 0.3512 | 0.2866 | 0.3029 | 0.2867 | 0.3024 | 0.2882 | 0.2626 | 0.2653 | 0.2748 | 0.2637 | 0.271 |
| Duba | 0.5915 | 0.6357 | 0.6709 | 0.6859 | 0.7114 | 0.7221 | 0.7151 | 0.4081 | 0.5592 | 0.4514 | 0.6198 | 0.6432 |
| Pseudot2 | 17.9566 | 17.7652 | 10.2991 | 5.952 | 2.4341 | 5.773 | 4.3831 | 5.8016 | 3.1524 | 6.0764 | 3.0674 | 4.437 |
| Beale | 5.91 | 4.9334 | 4.1601 | 3.778 | 3.09 | 3.2063 | 3.2458 | 10.3111 | 5.6028 | 8.9995 | 4.5431 | 4.3809 |
| Ratkowsky | 0.5197 | 0.4479 | 0.4078 | 0.3753 | 0.3489 | 0.3311 | 0.3131 | 0.2987 | 0.2854 | 0.2745 | 0.2645 | 0.2557 |
| Ball | 0.3693 | 0.1971 | 0.1134 | 0.0772 | 0.0573 | 0.0447 | 0.0354 | 0.029 | 0.0242 | 0.0205 | 0.0176 | 0.0153 |
| Ptbiserial | 0.6999 | 0.6354 | 0.563 | 0.5286 | 0.5021 | 0.5042 | 0.4609 | 0.4335 | 0.428 | 0.4235 | 0.4117 | 0.4013 |
| Frey | 2.0669 | 1.1674 | 0.7227 | 0.7507 | 0.1114 | 0.8524 | 0.6012 | 0.3979 | 0.3326 | 0.967 | 0.5389 | 1.1561 |
| McClain | 0.5019 | 0.7755 | 1.3095 | 1.6882 | 1.989 | 2.0047 | 2.5446 | 2.9865 | 3.0929 | 3.1872 | 3.3988 | 3.6109 |
| Dunn | 0.2354 | 0.2354 | 0.2247 | 0.2414 | 0.2772 | 0.305 | 0.3392 | 0.3534 | 0.3534 | 0.3534 | 0.3534 | 0.3534 |
| Hubert | 0.7243 | 0.8391 | 0.9718 | 0.9834 | 0.9826 | 0.9852 | 0.9858 | 1.0331 | 1.039 | 1.0384 | 1.0386 | 1.0393 |
| Sdindex | 17.4703 | 23.6931 | 23.3948 | 25.3816 | 28.4921 | 27.1434 | 31.6593 | 30.5158 | 30.2048 | 30.4642 | 31.2451 | 30.7231 |
| Dindex | 0.1051 | 0.0931 | 0.0815 | 0.0756 | 0.072 | 0.0688 | 0.0658 | 0.0631 | 0.0604 | 0.0583 | 0.0559 | 0.0542 |
| SDbw | 0.3663 | 0.2747 | 0.2154 | 0.1908 | 0.179 | 0.1394 | 0.1357 | 0.1324 | 0.1234 | 0.1138 | 0.103 | 0.0991 |

Supplemental Table 1. Clustering metrics across variety of clustering solutions ranging from 2 to 13 clusters.
